# ATP is a major determinant of phototrophic bacterial longevity in growth arrest

**DOI:** 10.1101/2023.01.04.522825

**Authors:** Liang Yin, Hongyu Ma, Elizabeth M. Fones, David R. Morris, Caroline S. Harwood

**Affiliations:** Department of Microbiology, University of Washington, Seattle, USA; Department of Environmental Science and Engineering, Xi’an Jiaotong University, Xi’an, P.R. China; Department of Biochemistry, University of Washington, Seattle, USA

**Author notes:** Corresponding author, Department of Microbiology, University of Washington, Box 357735, 1705 NE Pacific Street, Seattle, WA 98195-7735.

## Abstract

How bacteria transition into growth arrest as part of stationary phase has been well-studied, but our knowledge of features that help cells to stay alive in the following days and weeks is incomplete. Most studies have used heterotrophic bacteria that are growth-arrested by depletion of substrates used for both biosynthesis and energy generation, making is difficult to disentangle the effects of the two. In contrast, when grown anaerobically in light, the phototrophic bacterium *Rhodopseudomonas palustris* generates ATP from light via cyclic photophosphorylation and builds biomolecules from organic substrates such as acetate. As such, energy generation and carbon utilization are independent from one another. Here we compared the physiological and molecular responses of *R. palustris* to growth arrest caused by carbon source depletion in light (energy-replete) and dark (energy-depleted) conditions. Both sets of cells remained viable for six to ten days, at which point dark-incubated cells lost viability whereas light-incubated cells remained fully viable for 60 days. Dark-incubated cells were depleted in intracellular ATP prior to losing viability, suggesting that ATP depletion is a cause of cell death. Dark-incubated cells also shut down measurable protein synthesis, whereas light-incubated cells continued to synthesize proteins at low levels. Cells incubated in both conditions continued to transcribe genes. We suggest that *R. palustris* may completely shut down protein synthesis in dark, energy-depleted, conditions as a strategy to survive the nighttime hours of day/night cycles it experiences in nature, where there is a predictable source of energy in the form of sunlight during days.

**IMPORTANCE:** The molecular and physiological basis of bacterial longevity in growth arrest is important to investigate for several reasons. Such investigations could improve treatment of chronic infections, advance use of non-growing bacteria as biocatalysts to make high yields of value-added products, and improve estimates of microbial activities in natural habitats, where cells are often growing slowly or not at all. Here we compared survival of the phototrophic bacterium *Rhodopseudomonas palustris* under conditions where it generates ATP (incubation in light) and where it does not generate ATP (incubation in dark) to directly assess effects of energy depletion on long-term viability. We found that ATP is important for long-term survival over weeks. However, *R. palustris* survives 12h periods of ATP depletion without loss of viability, apparently in anticipation of sunrise and restoration of its ability to generate ATP. Our work suggests that cells respond to ATP depletion by shutting down protein synthesis.

## INTRODUCTION

Microorganisms are defined by their growth curves. Studies of model heterotrophic bacterial species like *E. coli* have taught us a great deal about how cells grow and how metabolism is reprogrammed as cells slow their growth rate and enter stationary phase (1). Less studied are strategies used by microbes to survive for long periods in growth arrest. Growth arrest can be caused by many factors including starvation for carbon, nitrogen, phosphate, or other nutrients that cells need for growth and replication. Many nutrients exist in growth-limiting amounts in natural environments (2), and bacteria can survive for long periods of time when growing very slowly or not at all (3–5). There are practical reasons to better understand the physiology of non-growing bacteria. In infectious disease, there is evidence that bacteria in growth arrest or those exhibiting very slow growth are more tolerant to antibiotics, and in the realm of biotechnology, growth-arrested bacteria are better biocatalysts because they can divert cellular resources to conversion of substrates to value-added products instead of to growth (6–8).

We have been exploring the molecular basis of bacterial longevity in growth arrest using the phototrophic alpha-proteobacterium *Rhodopseudomonas palustris* as a model. This microbe has extreme metabolic versatility (9, 10) and has received attention as a potential biotechnology chassis organism (11–13). An advantage that it has over heterotrophic bacteria for studies of longevity is that it generates all its ATP from light by photophosphorylation (Fig. 1). It derives carbon for biosynthesis from organic compounds but does not metabolize them for energy. This allows us to study growth arrest caused by nutrient limitation without the confounding effects of energy depletion that inevitably occur as heterotrophic cells struggle to stay alive when starved for an essential nutrient such as carbon. *R. palustris* cells that stop growing due to depletion of carbon or nitrogen remain viable for months when incubated in light (8, 14). Evidence that growth-arrested *R. palustri*s cells are not undergoing cycles of growth and death on long time scales include insensitivity to antibiotics that inhibit cell wall growth or DNA replication and experiments showing that such cells maintain unstable plasmids that are lost during cell division (15).

**Figure 1.**
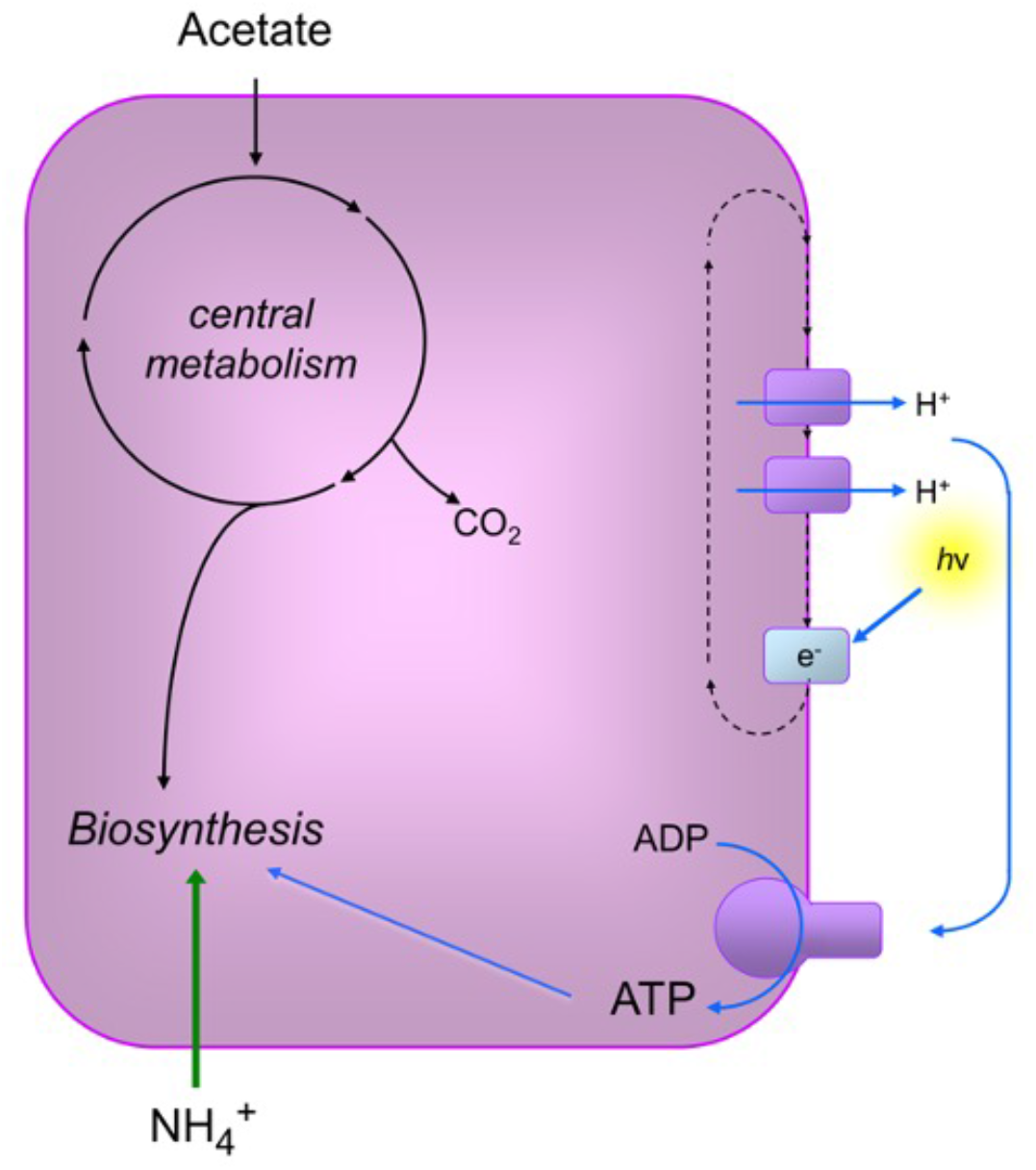
Diagram of *R. palustris* metabolism used to support growth prior to growth arrest. Cells in the experiments described here were grown photoheterotrophically with light (h*v*) as the energy source used by cells to generate ATP by cyclic photophosphorylation. Ammonium (NH_4_^+^) was provided in excess as a nitrogen source and acetate was supplied as the carbon source, which is not used in ATP-generating pathways and is used only to produce biomass. Cells were grown anaerobically. To achieve growth arrest, cells were given growth limiting amount of acetate, such that they became growth arrested when this carbon source was depleted. Growth-arrested cells continue to generate ATP from light but lose their ability to generate energy when incubated in continuous darkness.

We have hypothesized that the extraordinary longevity of *R. palustris* is related to its ability to generate ATP from light because we found that cultures die if moved to dark incubation conditions after entry into growth arrest (14). Here, we tested this hypothesis by comparing growth-arrested *R. palustris* cells incubated in light or dark conditions. We found that intracellular ATP levels correlated with both light availability and cell viabilities in growth arrest, with decreases in ATP preceding losses in viability. Both light and dark-incubated cells were transcriptionally active. However, dark-incubated cells were translationally inactive and formed hibernating ribosomes, whereas light-incubated cells continued to synthesize proteins. We suggest that a strategy of turning off and on protein synthesis may be an adaptation that *R. palustris* uses to survive the day-night cycles that it experiences in nature.

## RESULTS

### Conditions of growth arrest

*R*. *palustris* was grown anaerobically in light in sealed glass tubes containing mineral-salts medium until cells stopped growing due to depletion of the carbon source acetate (15). We assigned the time at which the optical density of cultures stopped increasing as “day 0” of growth arrest. To achieve dark-incubation conditions, tubes of growth-arrested cells were covered with aluminum foil or moved to dark incubators.

### Growth-arrested cells stay alive during long intervals of darkness, but not in continuous darkness

As shown in Fig. 2a, *R. palustris* cultures incubated in moderately bright light (40 μmol photons/m^2^/s; equivalent to light from a 60 W incandescent light bulb) maintained viability for a period of 25 d following growth arrest due to depletion of the carbon source acetate. The same was true for cells grown and incubated following growth arrest in dim light (4 μmol photons/m^2^/s; equivalent to the light from a 15 W incandescent light bulb). In nature, *R. palustris* is on a day-night cycle and we wondered how much darkness growth-arrested cells could tolerate in a 24 h period before losing viability. Non-growing cells exposed to continuous light, a 12 h light-12 h dark cycle or a 3 h light – 21 h dark cycle, remained fully viable for 60 d. Cells incubated with 1 h of light in a 24 h period or incubated in continuous darkness, started to lose viability about 6 d after growth arrest, and viabilities declined thereafter, with cells exposed to 1 h of light per 24 h losing viability less rapidly than cells incubated in continuous darkness (Fig. 2b).

**Figure 2.**
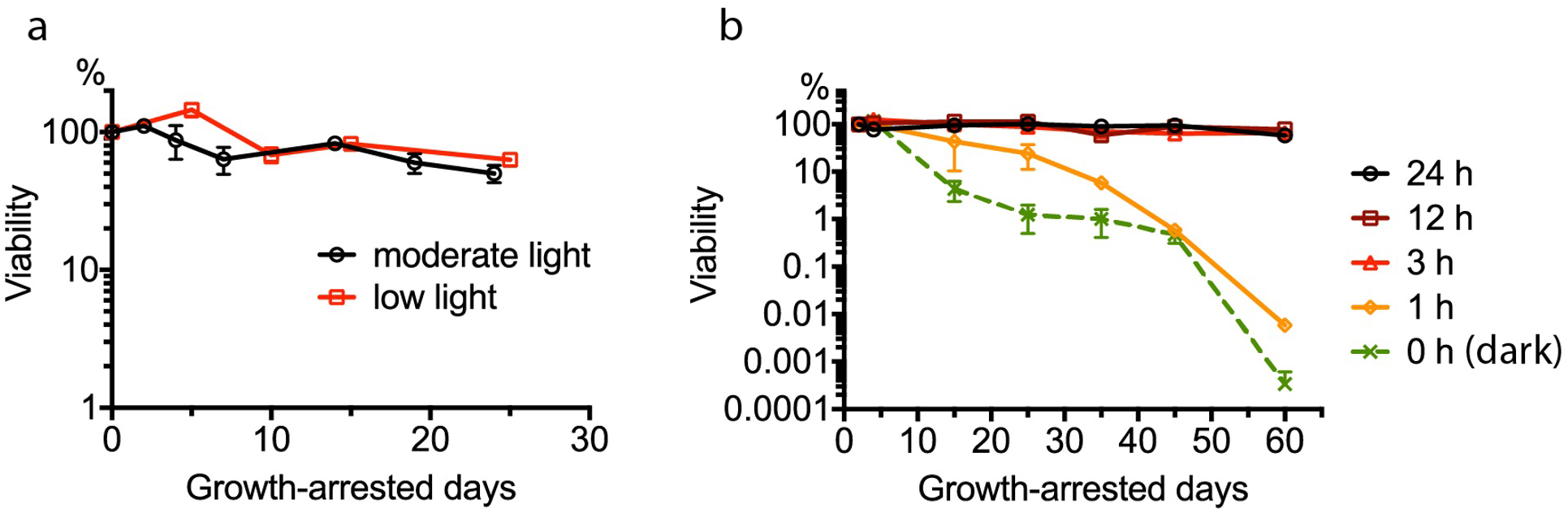
(a) Viability of *R. palustris* after growth arrest and incubation in moderate light (60W incandescent light bulb placed 10 cm away) or low light (15 W incandescent light bulb placed 10 cm away). Error bars represent standard deviations (n = 2) (b) Viability of *R. palustris* after growth arrest and incubation for variable amounts of time (in hours) in light per 24 h period. Cells were incubated in moderate light and moved to dark incubators for the dark periods of each 24 h day. Error bars represent standard deviations (n = 2)

### Dark-incubated growth-arrested cells become depleted in ATP

Intracellular ATP levels were approximately 15 nmol/mg protein when *R. palustris* entered growth arrest and dropped to about 7 nmol/mg total protein in cells incubated for 25 d in constant light (Fig. 3). In cells moved to dark incubation conditions immediately following growth arrest, ATP levels dropped to about 3.5 nmol/mg protein after 6 d, at which point cells were fully viable. At 25 d of dark incubation, intracellular ATP was below the limit of detection (about 0.1 nmol/mg protein) and this correlated with a three log decrease in viability (Fig. 3). Intracellular levels of ATP in non-growing cells incubated on 12 h light – 12 h dark cycles for a period of 25 d are shown in Fig. 4. We found that after 8 d and 25 d, ATP levels dropped to below the level of detection during the 12 h period of darkness but rebounded when cells were subsequently exposed to light.

**Figure 3.**
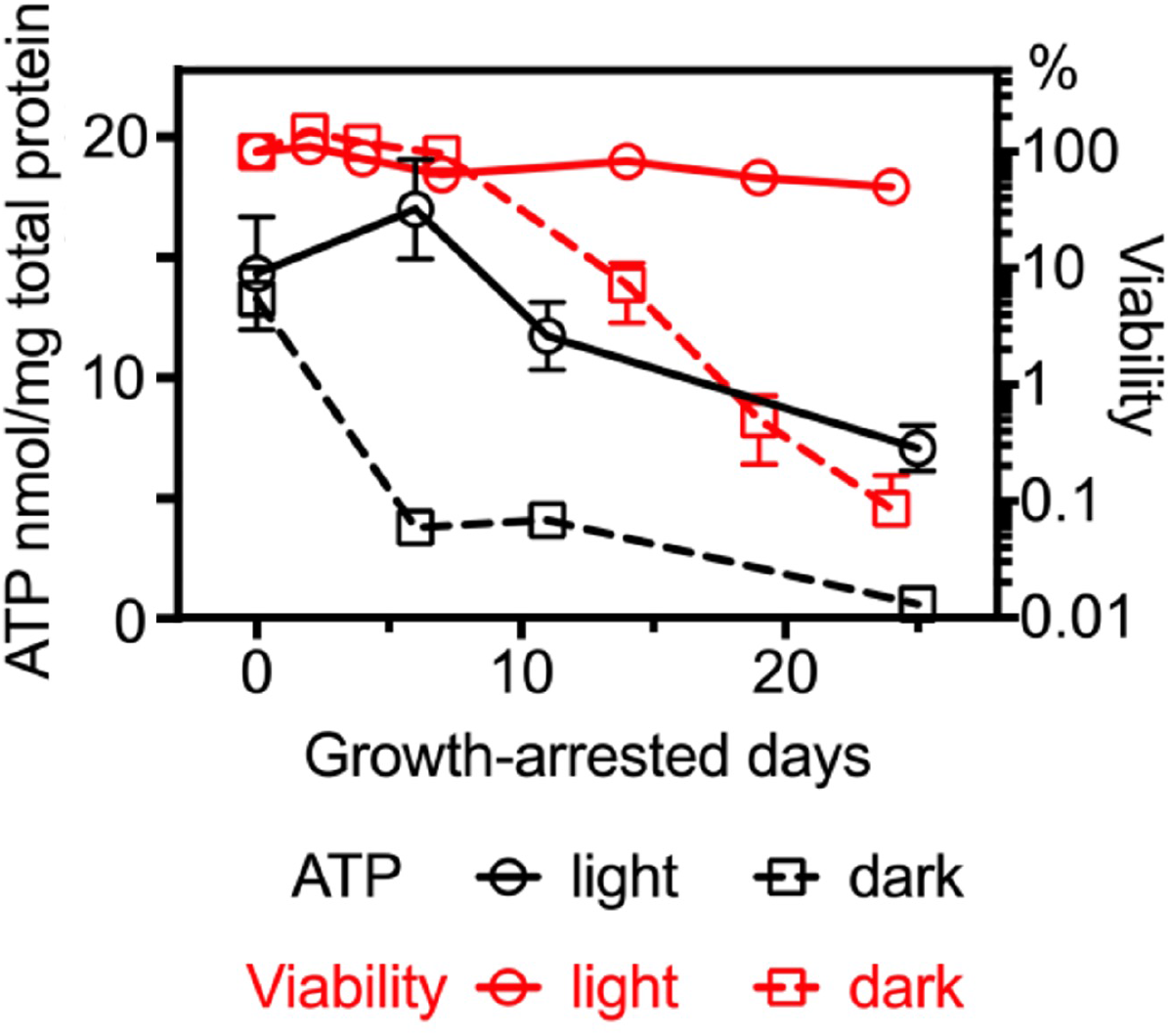
Viabilities and intracellular ATP content of *R. palustris* incubated in constant light or in constant dark following growth arrest. Error bars represent standard deviations (n=2).

**Figure 4.**
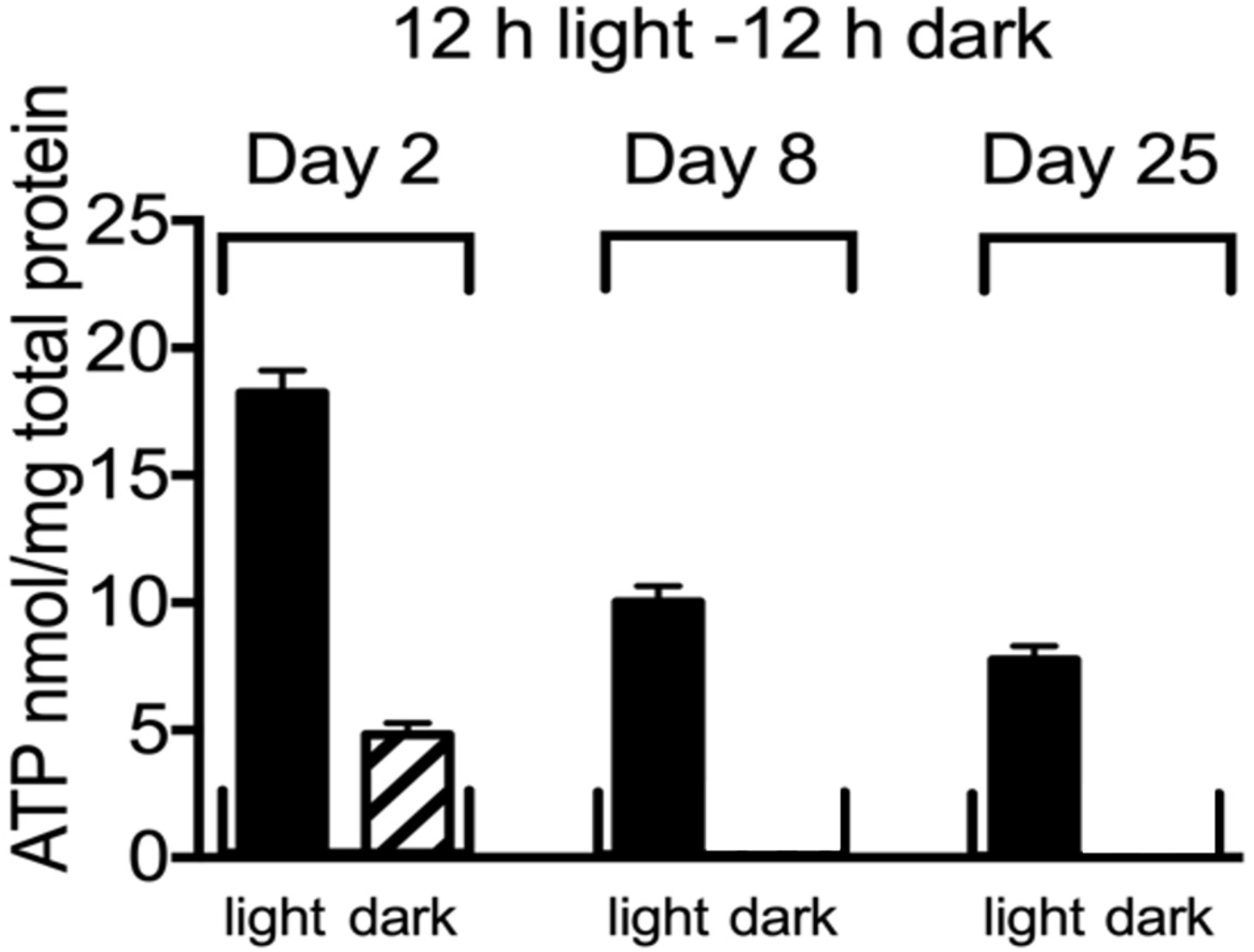
Intracellular ATP content of *R. palustris* in growth arrest day 0, day 8 and day 25 of continuous cycles of 12 h light - 12 h dark incubation. Error bars represent standard deviations (n=2).

### Non-growing cells incubated in dark shut down protein synthesis

We have previously reported that light-incubated *R. palustris* cells reduce but do not stop protein synthesis in growth arrest, and this continued protein synthesis is required for viability (15). By contrast, growth-arrested cells incubated in dark, did not appear to synthesize proteins. For example, cells incubated in dark and carrying an inducible *lacZ* gene *in trans*, did not synthesize significant amounts of LacZ protein at day 6 post-growth arrest, whereas light-incubated cells did synthesize LacZ (Fig. 5a). The inducer, phenylacetyl-homoserine lactone (PA-HSL) is diffusible across the cell membrane (16). We have previously shown that growth-arrested cells express substantial levels of LacZ in the absence of inducer, but levels are about 50% higher in the presence of PA-HSL (15). The ribosomes of growth-arrested cells incubated in light exist as populations of 30S, 50S and 70S species, which is similar to the ribosome subunit profile of growing cells (15). However, the ribosome profile of dark-incubated growth-arrested cells had one dominant peak of a 100S population of ribosomes (Fig. 5b). This 100S form has been widely observed in bacteria under nutrient starvation and is a translationally inactive dimer of two 70S ribosomes (17, 18). Another important aspect of translation is the charging states of tRNAs.

**Figure 5.**
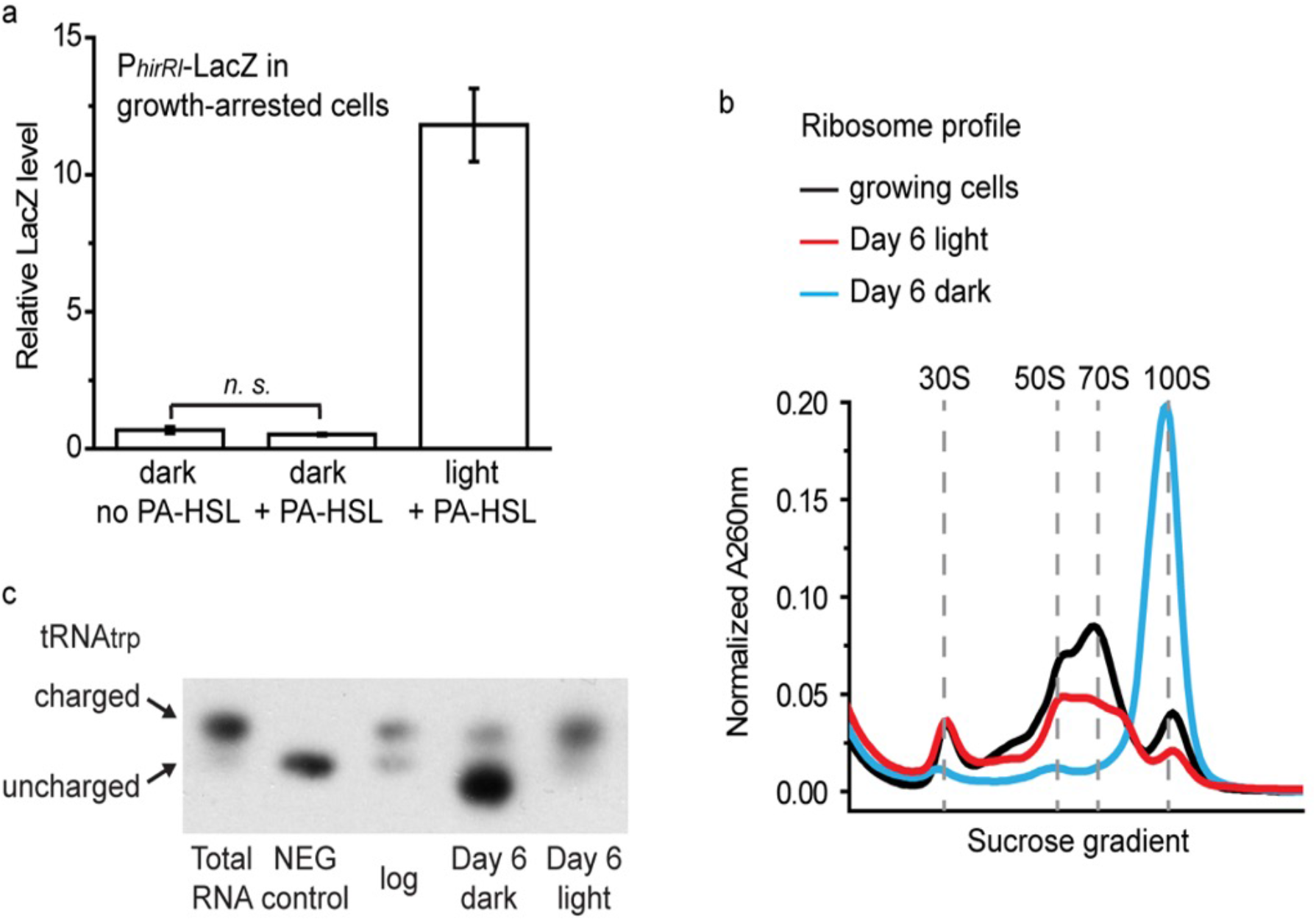
a) *R. palustris* carrying P_hirRI_-*lacZ in trans* on a plasmid was grown until growth arrest. The inducer PA-HSL (1 μM) was then added to cultures as indicated and cells were incubated as indicated. LacZ activity was measured after 2 d of incubation. b) Wild-type *R. palustris* was grown photoheterotrophically until acetate was depleted. The ribosome profile of light-incubated cells in growth arrest was reported previously (15) and is shown again here. For dark-incubated cells in growth arrest, ribosomes were purified and analyzed on a 7-47% sucrose gradient after 6 days incubation in dark. (c) The charging state of tRNA_trp_ in growing cells, or growth-arrested cells incubated in light or dark. A sample of uncharged tRNA was included as the negative control “NEG control,” as described previously (28).

During translation, tRNAs are aminoacylated (charged) with amino acids destined for delivery to ribosomes. The ratio of charged to free tRNAs is, therefore, crucial for the translation process. We have reported that about 60% of tRNA_trp_ is in the charged form, Trp-tRNA_trp_, in non-growing cells incubated in light (15). However, most of the tRNA_trp_, molecules were uncharged in dark-incubated non-growing cells (Fig. 5c). Taken together, these data indicate that growth-arrested *R. palustris* cells incubated in dark were not synthesizing measurable amounts of proteins.

### Dark-incubated, growth-arrested *R. palustris* cells are transcriptionally active

Even though growth-arrested cells incubated in dark were translationally inactive at day 6 post-growth arrest, RNA-seq experiments showed that they continued to synthesize mRNA. However, an analysis of the total number of sequencing reads generated from RNA Isolated from equivalent numbers of viable cells in exponential growth and at day 6 post growth arrest in light or in dark, showed that growth-arrested cells incubated under light or dark conditions both synthesized less RNA than exponentially growing cells and dark-incubated cells synthesized 35% less mRNA than light-incubated cells in growth arrest (SI Table S1). These differences, which are less than 2-fold, are not reflected in the RNA seq data presented in SI Tables S2 and S3, because calculation of RKPM (Reads Per Kilobase Million) values per gene and of fold changes between samples with statistical accuracy using DESeq2 both correct for differences in total sequence reads per sample. With this caveat, we found that about 500 genes were expressed at greater than 4-fold higher levels in dark- and light-incubated cells in growth arrest compared to cells in active growth and there was an overlap of about 200 genes between the two data sets (Fig. S1). The largest number of genes expressed at higher levels at day 6 post-growth arrest in both dark- and light-incubated cells as compared with exponentially growing cells fell into COG category S: function unknown (SI Fig S2). Dark-incubated cells had an enrichment of genes in category K, for transcription, that were expressed at higher levels.

Both dark- and light-incubated cells expressed about 700 genes at greater than 4-fold lower levels at day 6 post growth arrest compared to in active growth. Again, there was large overlap (∼450 genes) between the two data sets (SI Fig S1). For several categories of genes that were expressed at lower levels in both conditions, dark-incubated cells showed greater drops in gene expression than light-incubated cells (Fig. 6, SI Table S3). For example, 81% of the genes within the photosynthesis gene cluster (*rpa1505-1554*) that were down-regulated in both conditions showed greater decreases in gene expression in dark than in light. Similar patterns were observed for genes encoding ATP synthase subunits and ribosomal proteins (Fig. 6 and SI Table S3).

**Figure 6.**
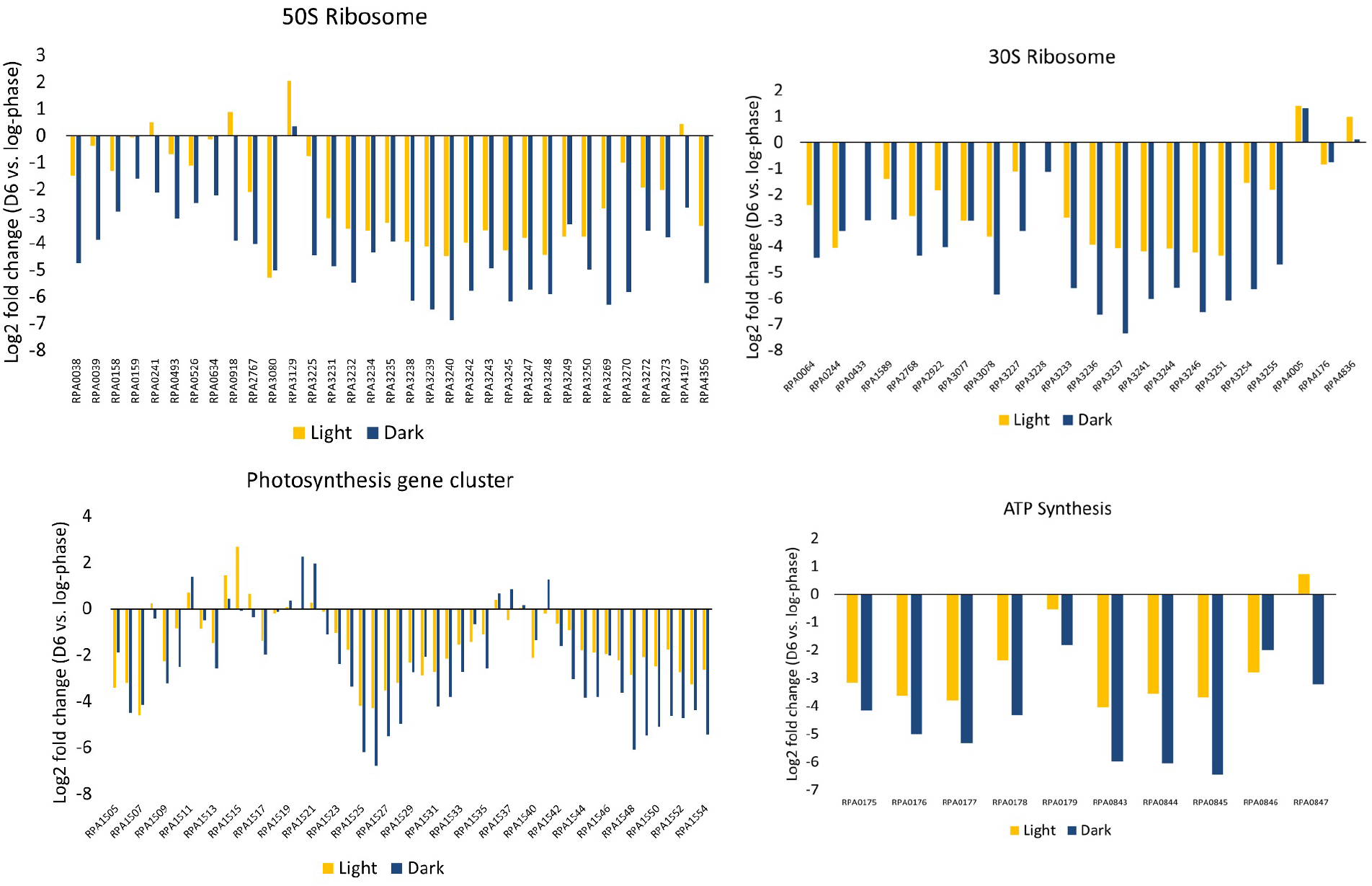
Comparison of fold-changes in expression of *R. palustris* photosynthesis genes, ribosomal protein genes and ATP synthase genes in light-incubated and dark-incubated cells at day 6 of growth arrest relative to growing cells.

As we have reported before (14), under illumination, most of the shift in transcription happens within the first day of growth arrest (SI Table S3). To examine if a longer period might induce additional changes, we measured the transcriptome at 20 d post-growth arrest in light but saw relatively few changes from day 6 to day 20. Of the 12 genes that showed a >4-fold increase in expression during this period, eight are predicted to encode hypothetical proteins with unknown function, one is a gene transfer agent (a phage-like entity) gene, two are transcriptional regulators, and one is a predicted permease (SI Table S3). The 38 genes that were expressed at lower levels at day 20 relative to day 6 post-growth arrest, included *cbbSL* genes encoding ribulose-bisphosphate carboxylase required for carbon dioxide fixation. These genes increased more than 15-fold in expression between day 1 and day 6 post-growth arrest, but then decreased in expression between day 6 and day 20. This may reflect a response to carbon dioxide that may be released from cells early in growth arrest as they metabolize carbon-storage compounds, like glycogen (19), but these sources of carbon dioxide diminish over time.

## DISCUSSION

Use of the phototropic bacterium *R. palustris* as a model to study longevity in growth arrest provides an opportunity to compare strategies that a bacterium uses to survive in energy-replete and energy-depleted situations. Cells in growth arrest due to carbon depletion maintained almost full viability for two months when incubated in light, but cells moved to dark immediately following growth arrest lost viability after an initial period of 6-10 days of full viability.

We found that although light-incubated cells maintained high levels of intracellular ATP over a period of 25 d following growth arrest, ATP levels dropped to undetectable levels over the same period in dark-incubated cells. These results are in line with those obtained with the same strain of *R. palustris* (CGA009) by Kanno et al.(20) In addition, these investigators measured the adenylate energy charge ([ATP] + 0.5 [ADP])/[ATP] + [ADP] + [AMP]) (21) of light and dark-incubated cells at day 5 post-growth arrest and found that whereas the energy charge of light-incubated cells matched that of growing cells, the energy charge of dark-incubated cells was dramatically lower. We note that intracellular ATP levels dropped in advance of losses of viability, suggesting that energy depletion caused cell death.

To identify consequences of ATP depletion we looked carefully at protein synthesis. Our rationale was that protein synthesis is an ATP-requiring process that can consume over 50% of the energy budget of bacteria (22). Dark-incubated cells appeared to carry out very little or no protein synthesis. Direct evidence for this is that dark-incubated cells did not translate a *lacZ* gene provided *in trans*. Indirect evidence is that the ribosome profile of dark-incubated cells consisted almost entirely of the 100S hibernating, translationally inactive form of ribosome and such cells did not have appreciable levels of amino-acylated tRNA_trp_.

Given that dark-incubated cells appeared not to be active in protein synthesis we were surprised to find that they continued to synthesize RNA at day 6 post growth arrest. Cells at this time point have about 20% of the amount of intracellular ATP that day 6 light-incubated cells have, and it may be that this is enough to support RNA synthesis. We did not identify genes known to be involved in RNA degradation or turnover to be upregulated in dark-incubated cells. Although we can’t exclude the possibility that dark-incubated cells maintain a subset of ribosomes that are translationally active, it is also possible that these cells have evolved to continue to synthesize RNA in anticipation that they will encounter light and quickly generate sufficient ATP to resume protein synthesis. This might explain why the transcriptional responses of both dark- and light-incubated cells in growth arrest are so similar. Differences in transcriptional responses in the two conditions could reflect a response of dark-incubated cells to prolonged ATP depletion and increasing physiological dysregulation that will result in cell death.

*R. palustris* resembles other bacteria in that it produces (p)ppGpp as it enters stationary phase, and this is essential for its longevity in growth arrest when illuminated (15). *E. coli* and other gram-negative bacteria respond to depletion of amino acids and other carbon and energy sources by synthesizing (p)ppGpp as part of the stringent response that includes decreased transcription of ribosomal protein genes and ribosomal RNAs, as well as down-modulation of ribosome maturation (1, 23). We note that the fold changes of downregulated ribosomal protein genes were much greater in dark-incubated cells than light-incubated cells, and perhaps a stringent response could explain this. It is known that the stringent response contributes to the ability of the cyanobacterium *Synechococcus elongatus* to adapt to darkness (24). However, the stringent response likely doesn’t explain all the transcriptional changes that we see and known transcription factors that regulate for example, expression of photosynthesis genes, were not altered in expression in growth-arrested cells incubated in either light or dark.

Gram-negative heterotrophic bacteria, including *Escherichia coli*, respond to growth arrest caused by energy and nutrient depletion by shrinking in size and there is evidence that they use their lipids, nucleic acids, and proteins as energy sources to maintain viability (1). *R. palustris* does not undergo a reduction in cell size during light or dark incubation following growth arrest, and although light and dark-incubated cells have markedly different metabolite profiles (20), there is no evidence that *R. palustris* CGA009 can generate ATP anaerobically in dark. We have reported that some level of protein synthesis is essential for longevity of growth-arrested *R. palustris* cells incubated in light and the same may be true for *E. coli*. In *E. coli*, hibernating ribosomes account for only about 60% of the total ribosome pool in stationary phase cells (25) and although *E. coli* has other mechanisms to shut down protein synthesis (26, 27), it is unclear whether it does so completely. Certainly, there is evidence that growth-arrested cells of *E. coli* and other gram-negative bacteria continue to synthesize proteins for days (4, 28). This suggests that some heterotrophic bacteria in growth arrest may resemble energy-replete growth-arrested *R. palustris* in prioritizing protein synthesis as a survival strategy.

If we extrapolate our observations of dark-incubated cells at 6 d post growth arrest to cells that have been incubated on a 12 h light-12 h dark cycle, the physiological response of *R. palustris* to dark makes sense. We found that even though *R. palustris* ATP levels were below the level of detection by day 8 in the dark phase of continuous 12 h light – 12 h dark cycles, intracellular ATP rebounded during the light phase. We hypothesize that *R. palustris* forms 100S ribosomes in dark as a way of preserving ribosomes that would then resolve to the 70S form in response to ATP generated from light. In short, *R. palustris* may have evolved to rapidly recommence protein synthesis in nature when ATP levels rise upon exposure to sun during the day. *R. palustris* has homologs of the cyanobacterial circadian clock genes *kaiB* and *kaiC* but is missing the *kaiA* clock gene (9). Circadian timekeeping in cyanobacteria is mediated by a phosphorylation-dephosphorylation cycle of KaiC that is driven by association and dissociation of a KaiA-KaiB-KaiC nanocomplex in a rhythmic cycle that maintains a self-sustained oscillation over 24 h periods of constant light (29). *R. palustris* has daily rhythms of KaiC phosphorylation in a regimen of 12 h light – 12 h dark, but the rhythm degenerates in constant light. Although *R. palustris* does not have true circadian rhythms as defined for cyanobacteria, a *kaiC* mutant has a growth defect when grown in 12 h light-12 h dark cycles but not when grown in continuous light (30). It will be interesting to test if this proto-circadian response may be involved in driving alternating periods of active and inactive protein synthesis in growth arrested cells.

## METHODS

### Bacterial strains, growth, and incubation conditions

*R. palustris* strain CGA009 was used as the wild type for this study as described before (15). Phototrophic cultures were grown anaerobically under illumination at 30°C in sealed glass tubes in defined PM medium (31) with 20 mM sodium acetate as the carbon source. When cultures reached their maximum optical density (approximately 2.0 × 10^9^ cells /ml), they were either maintained under illumination or moved to dark incubators to achieve dark incubation conditions. The viability of *R. palustris* cultures was determined by counting colony-forming units.

### ATP measurements

At desired time points, 0.9 ml culture was harvested by centrifugation. Cell pellets were immediately frozen in liquid nitrogen and stored at −80°C until used in ATP assays. For ATP measurements, frozen samples were resuspended with 1 ml 1% trichloroacetic acid and incubated for about 20 min at 30°C until cells were lysed. Cell lysate was centrifuged at maximum speed in a microcentrifuge for 2 min. The supernatant (about 500μl) was collected, and the pH of the supernatant was adjusted to ∼7.8 with 10M KOH and 25μl of 200mM Tris-Base. ATP in the supernatant was determined with an ATP bioluminescent assay kit according to instructions from the manufacturer (Sigma-Aldrich, FLAA-1KT).

### Ribosome purification and tRNA analysis

Ribosomes were purified and analyzed on a sucrose gradient as described previously (15, 32). The charging state of R. *palustris* transfer RNA was determined as described previously (31).

### LacZ assays

The inducible *lacZ* reporter was constructed as previously described and LacZ assays were as previously described (15). Briefly, *R. palustris* carrying P_*hirRI*_-*lacZ in trans* on a plasmid was grown until growth arrest. The inducer phenylacetate-homoserine lactone (PA-HSL) was then added to light- and dark-incubated cultures on day 6 of growth arrest. LacZ activity was measured after 2 d of incubation. The LacZ activity shown in Fig 5a is relative to the baseline of detectable activity in our assays.

### RNA-seq analysis

*R. palustris* was grown in PM medium as described above. Samples from the mid-logarithmic phase of growth were collected at OD_660_ = ∼0.6. After cultures stopped growing due to carbon depletion, samples were taken 1, 6 and 20 d post-growth arrest. Biological duplicates of samples were treated as follows. RNA was extracted from 5 ml samples using the miRNAeasy mini kit (Qiagen), treated with TURBO DNase (Ambion) and purified with RNeasy MinElute Cleanup kit (Qiagen). The samples were then sent to Genewiz, Inc. for library preparation and HiSeq RNA-seq sequencing. Raw RNA-seq reads were quality filtered and trimmed of adapters with Trimmomatic v0.39 (33) and the following parameter settings: HEADCROP:15 LEADING:3 TRAILING:3 SLIDINGWINDOW:4:15 MINLEN:35. Surviving read quality was assessed with FastQC (34). Reads were aligned to the *R. palustris* CGA009 reference genome and residual rRNA/tRNA reads were removed using Strand NGS v4.0, build 242089 (© Strand Life Sciences, Bangalore, India) following default parameters. Differential gene expression analysis was performed using DESeq2 (fold-change ≥ 4, *p* < 0.05)(35). Sequencing results were processed and analyzed in house with StrandNGS (strand-ngs.com).

## DATA AVAILBILITY

All data are supplied as supplementary files. Raw data are available on request from the corresponding author.

## ACKNOWLEDGMENTS

This work was supported by the US Army Research Office, contract W911NF2110015.

## SUPPLEMENTAL TABLE AND FIGURE LEGENDS

Figure S1. Genes increased and decreased in expression four-fold or more relative to growing cells after 6 d incubation in light or dark following growth arrest.

Figure S2. Transcript profile of *R. palustris* cells incubated in light or dark for 6 days after onset of growth arrest. Genes whose expression changed significantly (>4-fold, p<0.05) from log-phase growth were classified by NCBI Clusters of Orthologous Genes (COG). COG Categories: C-Energy production/conversion, D-Cell cycle control, E-Amino acid transport/metabolism, F-Nucleotide transport/metabolism, G-Carbohydrate transport/metabolism, H-Coenzyme transport/metabolism, I-Lipid transport/metabolism, J-Translation/ribosomal structure and biogenesis, K-Transcription, L-Replication, M-Cell wall/membrane/envelope biosynthesis, N-Cell motility, O-Post-translational modification, P-Inorganic ion transport/metabolism, Q-Secondary metabolites biosynthesis, S-Unknown, T-Signal transduction mechanisms, U-Intracellular trafficking, V-Defense mechanisms, None-No COG assigned.

SI Table S1. Number of reads generated by RNA sequencing before and after quality-filtering and alignment to the reference transcriptome.

SI Table S2. RPKM (Reads Per Kilobase Million) of RNA isolated from *R. palustris* in log-phase or in growth arrest incubated in dark or in light for the number of days indicated. D1, D6, and D20 refer to day 1, day 6, and day 20 of growth arrest, respectively.

SI Table S3. List of genes that are differentially expressed (>= 2-fold change, p < 0.05) between the experimental conditions described in each tab with RPKM values that exceeded 50 for at least one of the conditions described.

